# Assembly of whole-chromosome pseudomolecules for polyploid plant genomes using outcrossed mapping populations

**DOI:** 10.1101/119271

**Authors:** Chenxi Zhou, Bode Olukolu, Dorcus C. Gemenet, Shan Wu, Wolfgang Gruneberg, Minh Duc Cao, Zhangjun Fei, Zhao-Bang Zeng, Andrew W. George, Awais Khan, G. Craig Yencho, Lachlan J.M. Coin

**Affiliations:** Institute for Molecular Bioscience, University of Queensland, St Lucia, Brisbane, QLD 4072, Australia; Department of Horticulture, North Carolina State University, Raleigh 27695, USA; International Potato Center, PO. Box 1558, Lima 12, Peru; Boyce Thompson Institute, Cornell University, Ithaca, NY 14853, USA; Department of Statistics and Bioinformatics Research Center, North Carolina State University, Raleigh 27695, USA; Data61, CSIRO, Ecosciences Precinct, Brisbane, QLD 4102, Australia; Department of Plant Pathology and Plant-microbe Biology, Cornell University, Geneva, NY 14456, USA

## Abstract

The assembly of whole-chromosome pseudomolecules for plant genomes remains challenging due to polyploidy and high repeat content. We developed an approach for constructing complete pseudomolecules for polyploid species using genotyping-by-sequencing data from outcrossing mapping populations coupled with high coverage whole genome sequence data of a reference genome. Our approach combines *de novo* assembly with linkage mapping to arrange scaffolds into pseudomolecules. We show that the method is able to reconstruct simulated chromosomes for both diploid and tetraploid genomes. Comparisons to three existing genetic mapping tools show that our method outperforms the other methods in accuracy on both grouping and ordering, and is robust to the presence of substantial amounts of missing data and genotyping errors. We applied our method to three real datasets including a diploid *Ipomoea trifida* and two tetraploid potato mapping populations. The linkage maps show significant concordance with the reference chromosomes. We resolved seven assembly errors for the published *Ipomoea trifida* genome assembly as well as anchored an unplaced scaffold in the published potato genome.

## Introduction

High quality genome assembly plays an essential role in plant genomic and genetic analyses. The construction of a genome assembly typically adopts a ‘bottom-up’ architecture. Short sequencing reads are first assembled by analysing read overlaps to build contigs^1^. Contigs are then bridged to construct scaffolds using long reads or large insert size paired reads^2^. This process is sometimes repeated for multiple rounds by gradually introducing larger insert size libraries. Finally, long distance information is integrated to order scaffolds to establish pseudomolecules^3^. Several tools have been proposed for genome assembly along these lines^4, 5^. However, these tools are often limited in polyploid plant genomes due to high levels of heterozygosity, the large amount of repetitive DNA, as well as increased complexity in resolving haplotypes which scales exponentially in the number of homologous chromsomes.

Chromosome-scale scaffolding using long distance information is a crucial step in generating high quality genome assemblies. A variety of mapping information could be utilised, such as physical maps^6^, genetic maps^7^, optical maps^8^, syntenic maps^9^ and chromatin interactions^3^. Genetic mapping has been widely adopted as it generates longer range information than physical mapping techniques. The most commonly used bacterial artificial chromosome (BAC) approach generally handles insert sizes up to 350kb^10^, which could be too short to flank heterochromatin or long repetitive regions^11^. In comparison, theoretically, genetic mapping provides linkage information as long as two contigs are located on the same chromosome.

Chromosome-scale linkage analysis requires high marker density in mapping population. In order to obtain the full map of the chromosomes, markers should cover as much of the genome as possible. Genotyping-by-sequencing (GBS) provides a cost-effective approach which enables large-scale detection of biomarkers at low coverage with high missing and genotyping error rates^12^. Despite a reduced representation, usage of restriction enzyme ensures nearly even distribution of the markers across the whole genome. Barcoding systems make GBS highly multiplexed and suitable for genotyping a mapping population.

Linkage analysis for large scale marker sets requires high performance genetic mapping tools. Conventional tools such as MAPMAKER^13^ and Map Manager QTX^14^, and more recent tools such as R/qtl^15^ and AntMap^16^ have been optimised for relatively small but high quality marker sets thus can seldom process the tens of thousands of markers with high missing and error rates generated by GBS. Moreover, these tools were designed for inbred lines and cannot be applied to outcrossing mapping populations. Development of inbred lines, however, could be difficult, expensive or time-consuming, especially for polyploids^17^. Several methods have since been proposed for outbred lines^17–19^. However, none of them have been designed for polyploids. So far, most polyploid genetic linkage maps have been built using diploid models^20^. This limits markers that can be used for linkage analysis to a few specific segregation patterns compatible with diploid models, such as simplex and duplex markers^21, 22^.

Here we describe a novel method for constructing genetic linkage maps. The method relies on the availability of a high density marker set on a F1 outcrossed population and reference contigs or scaffolds. We focus on building genetic maps for marker blocks, rather than individual markers. Key features of this method include (1) it is accurate, (2) it is computationally effective, (3) it uses outcrossed mapping populations, (4) it is intrinsically suitable for polyploid species, and (5) it is robust for missing data and genotyping errors and (6) it detects assembly errors. Combined, these features enable us to build high quality pseudomolecules covering a large proportion of polyploid plant genomes. Using both simulated and real diploid and tetraploid datasets, we demonstrate substantial improvements of our approach over existing genetic mapping algorithms.

## Results

### Overview of method

We have developed a new method called PolyGembler (**Poly**ploid **Ge**netic-linkage ass**embler**) for assembly of polyploid genomes using genetic linkage information. Figure 1 provides an overview of PolyGembler, with details in the Online Methods. The method assumes availability of genome-wide genotyping data such as GBS and array data, collected on a F1 outbred family, as well as high coverage (i.e. greater than 30X) whole genome sequence data on a reference sample, or alternatively the availability of a set of reference contigs or scaffolds. Our approach combines *de novo* assembly with linkage mapping to arrange scaffolds into pseudomolecules. By mapping marker set to scaffolds we are able to infer scaffold haplotypes for each sample even in the presence of substantial amounts of missing data and genotyping errors. We use these haplotypes to infer linkage groups corresponding to chromosomes as well as the optimal ordering of scaffolds within these chromosomes. PolyGembler consists of three major modules, namely variant detection (Fig. 1a), recombination frequency (RF) estimation (Fig. 1b-d) and genetic mapping (Fig. 1e-f). The initial step is to use existing assembly graph algorithms to infer reference genome scaffolds. The variant detection module aligns GBS data to reference scaffolds to call SNPs. The RF estimation module infers haplotypes for each scaffold and then uses these haplotypes to detect assembly errors and calculate RFs between all pairs of scaffolds. Haplotyping accuracy can be improved by combining information from nearest-neighbour scaffolds (Online Methods). The genetic mapping module follows the conventional linkage map construction framework to build linkage groups of scaffolds and optimise the order of these scaffolds, using a modified traveling salesman problem (TSP). The scaffold-based genetic linkage maps are finally used to construct pseudomolecules.

**Figure 1.**
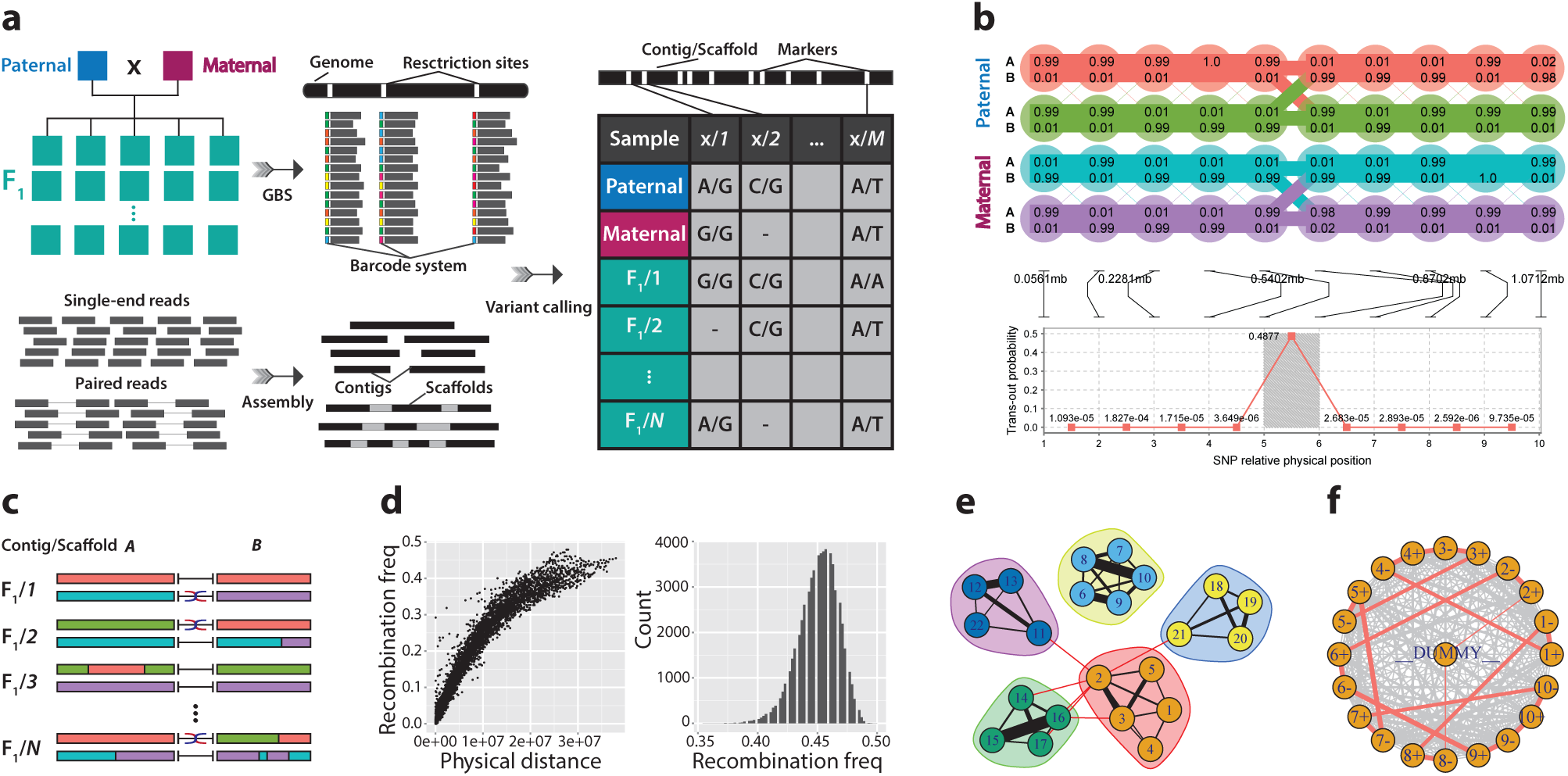
Polyploid genetic-linkage assembler (PolyGembler) framework. (a) GBS technology is used to sequence an outcrossed mapping population. High coverage whole genome sequence data is used to build reference contigs and scaffolds. By mapping GBS reads to the reference assembly, we call variants for the mapping population. (b) Construction of haplotypes for each scaffold and for each sample with a hidden Markov model (HMM). A well trained HMM for a diploid is depicted in the top panel. Here the scaffold consists of ten markers illustrated as ellipses. The numbers within ellipses indicate the probabilities of the parental haplotypes pass *A*- and *B*-allele. The width of the line connecting two ellipses is proportional to the RF between parental haplotypes at that position. Recombinations are prohibited between paternal and maternal haplotypes. The model assumes each F1 progeny inherits one haplotype from each parent’s gamete and maximises the likehood of the marker set. This model is able to identify assembly errors. As we can see, the RF between the 5th and 6th marker is abnormally large as described in the bottom panel. Therefore, it is highly likely an misassembly. In the middle panel, we show the positions of ten markers along the scaffold. (c) RF estimation between each pair of scaffolds by counting the mismatches of parental haplotypes. It should be noted here that, RFs are calculated at all four possible connection directions. Moreover, as haplotypes are constructed independently, we consider all possible correspondences between parental haplotypes. (d) The estimated RFs reflect the physical distances. Plots are generated from a simulated dataset. The dot plot and histogram is for scaffold pairs from the same chromosome and different chromosomes, respectively. (e) Linkage groups construction for the reference scaffolds. We build a graph for the scaffolds which is weighted by the estimated RFs. The linkage groups are then identified by a graph partitioning algorithm. (f) Ordering for each linkage group by solving a modified traveling salesman problem (TSP). The TSP treats the two ends of a scaffold as different nodes and is designed to ensure that the two ends are neighbours in the optimal solution. The dummy node, which has equal distances to all the other nodes, is introduced to convert a Hamiltonian circuit to a Hamiltonian path.

### Application to simulated Ipomoea trifida GBS data

We simulated a reference genome from the *Ipomoea trifida* sequencing data provided in^23^ to make it comparable to the real genome (Online Methods). The resulted genome consists of 15 chromosomes (Supplementary Table S3.). The total size is ~ 482Mb, of which, ~ 24% are repeated sequences. Based on the reference genome, we simulated GBS data for a diploid and a autotetraploid outcrossed F1 mapping population of 192 samples. In order to simulate missing data and genotyping errors, for any enzyme recognition site, the depth of sequencing was sampled from the truncated normal distribution *N*(5,5^2^). Under this assumption, approximately 15.87% recognition sites were missed with depth ≤ 0. We simulated autotetraploid mapping population twice. The resulted depth of coverage is 10.8× and 43.2× for diploid and autotetraploid simulation, respectively (Supplementary Note S1., Online Methods). We simulated a genome assembly from the reference genome (Online Methods). The total size is ~ 527Mb and the N50 statistic is ~ 106kb (Supplementary Table S5.). The genome assembly was used as the reference for variant calling from the GBS data.

#### Diploid simulation

The variant calling module identified 53,194 SNPs for pseudomolecule construction. These SNPs are distributed across 3,482 scaffolds. These scaffolds summed to ~ 394Mb cover ~ 82% of the genome. In order to investigate the accuracy of our method, we mapped these scaffolds to reference chromosomes to decide the true linkage groups, as well as physical distances between all pairs of scaffolds that belong to the same chromosome (Online Methods). 3,004 scaffolds were considered successfully mapped. For scaffold pairs that located on the same chromosome, the estimated RFs correlate with the physical distances very well (Fig. 2a). For scaffold pairs that mapped to different chromosomes, the RF estimations are consistently large and greater than the predefined threshold used for grouping (Fig. 2b). The linkage groups constructed by the method perfectly recognized 15 reference chromosomes. Within each linkage group, the order of the mapped scaffolds demonstrates a high correlation with the true order on the corresponding reference chromosome. Precisely the Spearman’s rank correlation coefficients between them ranged from 0.82 to 0.95. The estimated RFs and LOD scores between scaffolds clearly reflect the linkage information (Supplementary Figure S2.).The pseudomolecules are in concordance with the reference chromosomes with a few incorrect ordering at the end (Fig. 3a).

**Figure 2.**
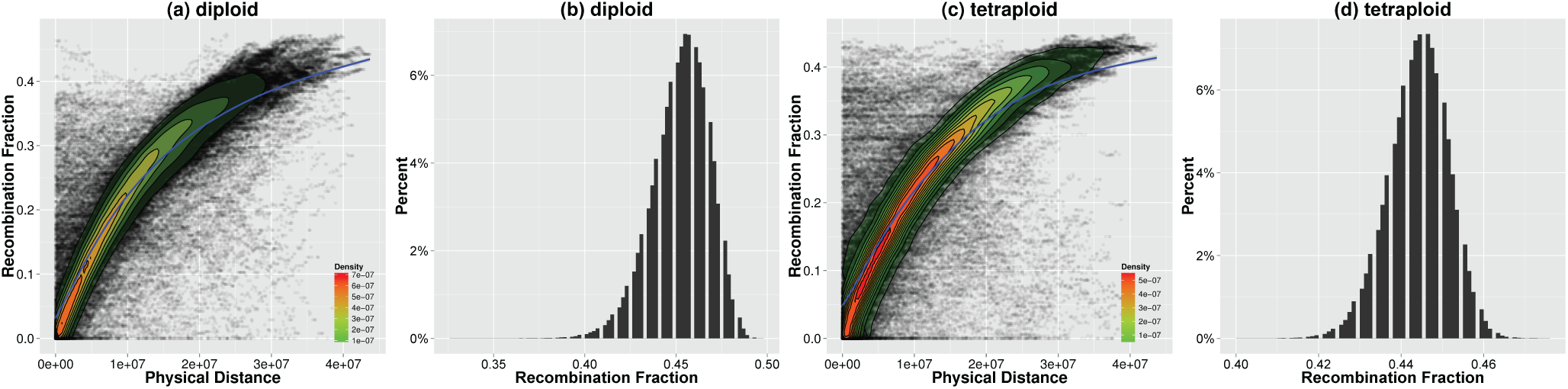
Estimated RFs versus physical distances plots for simulated data. (a) and (c), dot plots for the physical distances against the estimated RFs for scaffold pairs that mapped to the same reference chromosome. (b) and (d), histograms of the estimated RFs for scaffold pairs that mapped to different chromosomes. We added density plots (range from green to red) as well as smoothed conditional mean plots (blue) to dot plots. The smoothed conditional means were calculated using generalized additive model (GAM).

**Figure 3.**
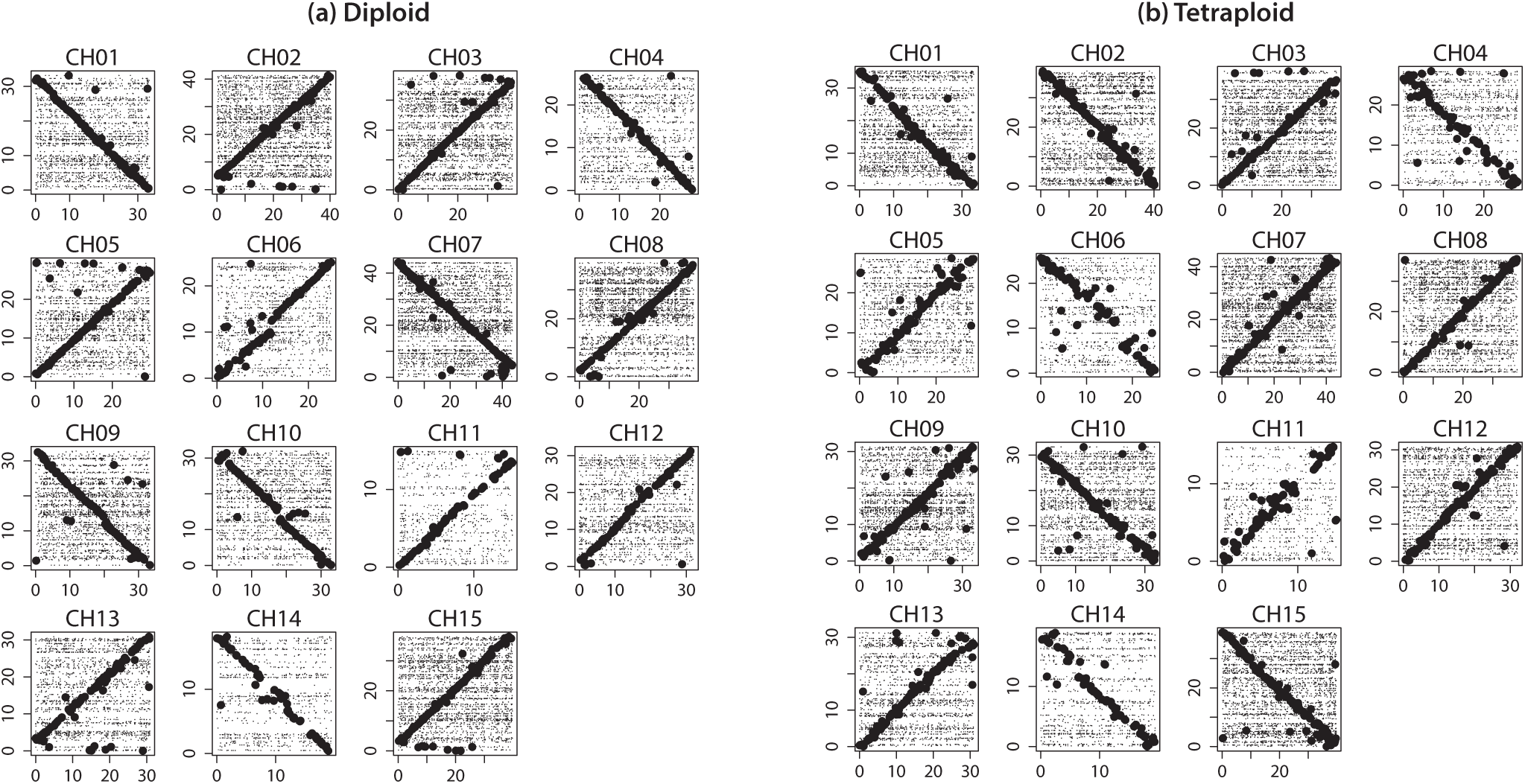
Collinear plots between the reference chromosomes and pseudomolecules constructed by the proposed method for simulated data. (a) diploid simulation and (b) tetraploid simulation. *x*- and *y*-axis represent the position (Mb) on reference chromosome and pseudomolecules, respectively. Each line in the plots represents a collinear alignment between the simulated reference chromosomes and the pseudomolecules that we built. Long collinear alignments (≥ 10Kb) are plotted with thick lines. As we can see, long collinear alignments form a diagonal line in each plot. This indicates the high correlations between the simulated reference chromosomes and the pseudomolecules built from the scaffolds. Two types of diagonal lines are observed, namely from the bottom-left corner to the top-right corner and from the top-left corner to the bottom-right corner. A diagonal line of the first type suggests that the scaffolds are arranged in the same order on the pseudomelecule as that on the simulated reference chromosome, compared to a reverse order arrangement of scaffolds suggested by a diagonal line of the second type.

#### Tetraploid simulation

The variant calling module identified 51,656 SNPs located on 3,435 scaffolds for pseudomolecule construction. These scaffolds summed to ~ 392Mb cover ~ 81% of the genome. 2,968 scaffolds were considered successfully mapped to the reference chromosomes. The RF estimations are similar to the diploid simulation (Fig. 2c and d, Supplementary Figure S2.). 15 pseudomolecules corresponding to reference chromosomes were constructed without miss grouping of the mapped scaffolds. The ordering is not as precise as that in the diploid simulation. The Spearman’s rank correlation coefficient was observed in the range from 0.81 to 0.94. The pseudomolecules still correlate highly with reference chromosomes (Fig. 3b). However, more incorrect ordering is observed especially in CH04, CH06, CH11 and CH14. There are two major reasons that cause the loss of accuracy for the pseudomolecules. Firstly, genotype calling for tetraploid genome is difficult. The accuracy of haplotype reconstruction is sacrificed to deal with the high genotyping error rates. Secondly, the haplotype accuracy was compromised by low levels of genetic diversity of the marker set. As the number of SNPs on a scaffold could be small, if many of them display low level of genetic diversities, the program can hardly distinguish parental haplotypes from each other, and might eventually lead to incorrect assignment of parental haplotypes to F1 progeny. In order to investigate the proposed method thoroughly, we also simulated GBS data at 20x. The accuracy of the RF estimations decreased. The linkage groups constructed by the method are still correct. The order of scaffolds within each linkage group, however, lost some accuracy (Supplementary Figure S3.).

### Application to real Ipomoea trifida GBS data

The *Ipomoea trifida* mapping population consists of 210 F1 progeny and two parents. It was derived from the crossing of two diploid (2n=2x=30) *Ipomoea trifida* heterozygous genotypes, called M9 × M19. The population was developed by the international potato center (CIP) and the DNA from the population was provided to NCSU for GBS. We used the *de novo* genome assembly ITR_r1.0^23^ as the reference for variant detection. The ITR_r1.0 genome assembly consists of 77,400 scaffolds. The total size is ~ 512Mb and the N50 statistic is ~ 43Kb. The variant detection module called 48,202 SNPs located on 2,282 scaffolds. These scaffolds summed to ~ 226Mb cover approximately 44% of the whole genome. As the coverage is small, it is difficult to estimate the gap size between the contiguous scaffolds. Therefore, instead of building pseudomolecules, we only constructed genetic linkage maps from the scaffolds. We used another *Ipomoea trifida* genome assembly NSP306v2 to validate the genetic linkage maps. NSP306v2 was assembled by the GT4SP project (http://sweetpotato.plantbiology.msu.edu/gt4sp_download.shtml). It consists of 30,377 scaffolds, and the N50 statistic is ~1.5Mb.

We detected seven misassemblies in ITR_r1.0 (Online Methods). Figure 4a depicts the RF estimations along the scaffold 15 based on 30 independent runs of the haplotype phasing algorithm (see Supplementary Figure S4. for the other six). As we can see, there is a potential misassembly site around 177kb, where the estimated RF is 0.321 ±0.117. By mapping it to NSP306v2 scaffolds, we found that while the first ~ 166kb was mapped to scaffold 4, the last ~ 138kb was mapped to scaffold 40002 (Fig. 4b and c). This result supports the misassembly of scaffold 15 as well as the misassembly position. In fact, 5/7 detected misassemblies were confirmed by mapping to NSP306v2 scaffolds (Supplementary Table S6.). We split the incorrect ITR_r1.0 scaffolds to generate new scaffolds and excluded the original ones from linkage analysis (Supplementary Table S6.). As a result, 2289 scaffolds were included in linkage map construction. The haplotype phasing algorithm failed to give any feasible results for 56 scaffolds. The remaining 2,233 scaffolds were assigned to 15 linkage groups (Supplementary Figure S2.). The total length of the genetic linkage maps is approximately 4058cM (Supplementary Figure S5.). Among the 2,233 scaffolds, 1,572 scaffolds were considered successfully mapped to NSP306v2 scaffolds thus could decide the linkage groups, order and pairwise distances (Online Methods).

**Figure 4.**
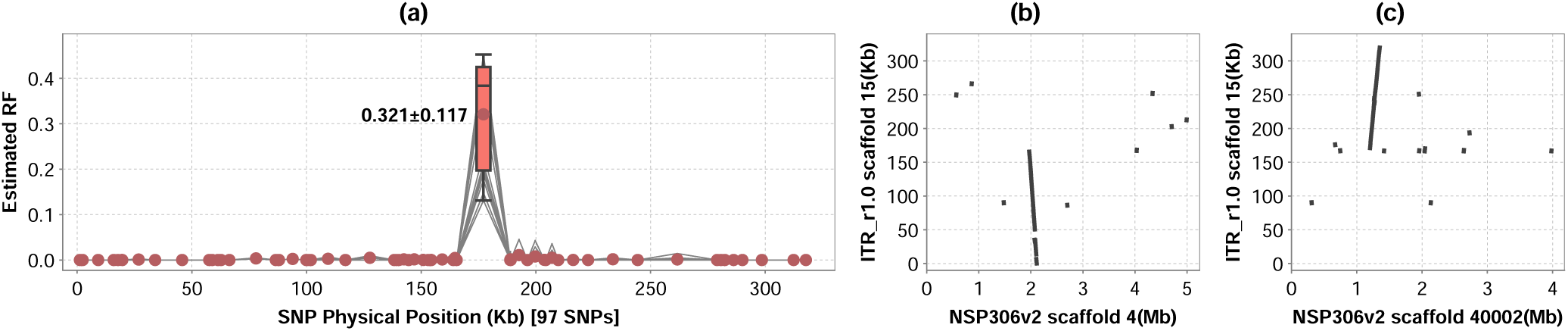
ITR_r1.0 scaffold 15 is a misassembly. (a) the estimated RFs along the scaffold based on 30 independent runs. Each gray line plot represents a independent run. *x*- and *y*-axis represent the SNP positions (Kb) on the scaffold 15 and the estimated RFs between the adjoining SNPs, respectively. Solid dots indicate the mean RFs at that position. There is a misassembly around 177Kb, where the estimated RFs are 0.321 ± 0.117. (b) and (c), the first ~ 166kb of scaffold 15 is collinear with NSP306v2 scaffold 4, while the last ~138kb is collinear with NSP306v2 scaffold 40002.

Figure 5a illustrates RF estimations for mapped scaffolds. There is a dot cluster at the top of the plot which represents overestimation of RFs. This is caused by discrepancies between assembly NSP306v2 and ITR_r1.0. According to the genetic linkage maps we built, 26 discrepancies were detected where ITR_r1.0 scaffolds that mapped to the same NSP306v2 scaffold were assigned to multiple linkage groups. For example, 119 ITR_r1.0 scaffolds were mapped to the NSP306v2 scaffold 1, however, were assigned to three linkage groups. We identified ten ≥ 1Mb misassemblies using BioNano maps in NSP306v2 (Supplementary Table S7.). All of them were identified as discrepancies by our method. The break points of the misassemblies agree with the BioNano mapping results. It is difficult to tell if the remaining 16 discrepancies we detected are true or false positives. Firstly, there might be genome structure variations between the two *Ipomoea trifida* lines that used for genome assembly. Secondly, the *Ipomoea trifida* genome abounds with repetitive sequences^23^ and makes the mapping process difficult. In the current study, an ITR_r1.0 scaffold is regarded as mapped if no less than 50% base pairs are collinear with a NSP306v2 scaffold (Online Methods). The low threshold could introduce false positive mapping. Indeed, if we increased the threshold to 70%, the number of mapped ITR_r1.0 scaffolds would decrease to 1200, and the number of discrepancies would decrease to 14. The overestimations disappeared after removing the discrepancies (Fig. 5b).

**Figure 5.**
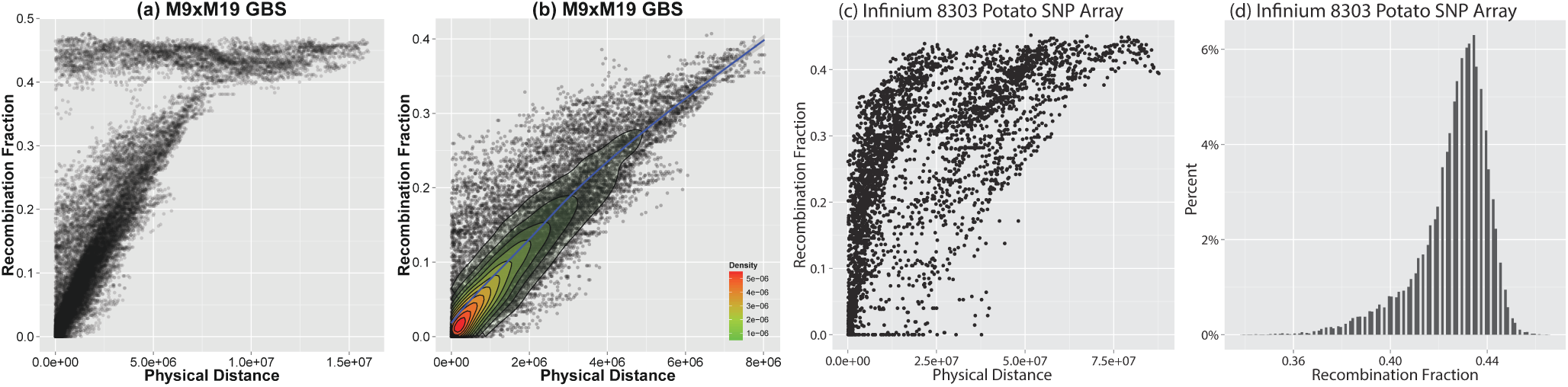
Estimated RFs versus physical distances plots for real data. (a) dot plots for the physical distance against the estimated RFs for ITR_r1.0 scaffold pairs that mapped to the same NSP306v2 scaffolds. (b) the discrepancies between NSP306v2 and ITR_r1.0 were removed. We added a density plot (range from green to red) as well as a smoothed conditional mean plot (blue) calculated using generalized additive model (GAM). (c) dot plots for the physical distance against the estimated RFs for scaffold pairs that come from the same pseudomolecule. (d) histograms of the estimated RFs for scaffold pairs that come from different pseudomolecules.

Figure 6 illustrates the syntenic maps between the genetic linkage maps we built and the NSP306v2 scaffolds. Here we only show three linkage groups that contain scaffolds mapped to NSP306v2 scaffold 1, and see Supplementary Figure S6. for the remaining 12 genetic linkage maps. The break points of scaffold 1 agree with those estimated from BioNano mapping (Supplementary Table S7.). In each linkage group, the order of the ITR_r1.0 scaffolds is almost identical to the mapping order on the scaffold 1. This is also true for the other scaffolds although with a few exceptions. NSP306v2 scaffold 48 is observed twice (Fig. 6a and c). Precisely, in the first linkage group, there is only one ITR_r1.0 scaffold that was mapped to scaffold 48, namely scaffold 167. Interestingly, this scaffold was also successfully mapped to NSP306v2 scaffold 40007. Precisely, this scaffold is approximately 146Kb long. While the first 57Kp was mapped to scaffold 40007, the last 89Kp was mapped to scaffold 48. As we chose the longest alignment, we considered it mapped to the later one. However, according to the linkage analysis, it is closer to those scaffolds that mapped to scaffold 40007. This is either because of the misassembly of scaffold 167 which has not been detected or genome diversity between two *Ipomoea trifida* lines.

**Figure 6.**
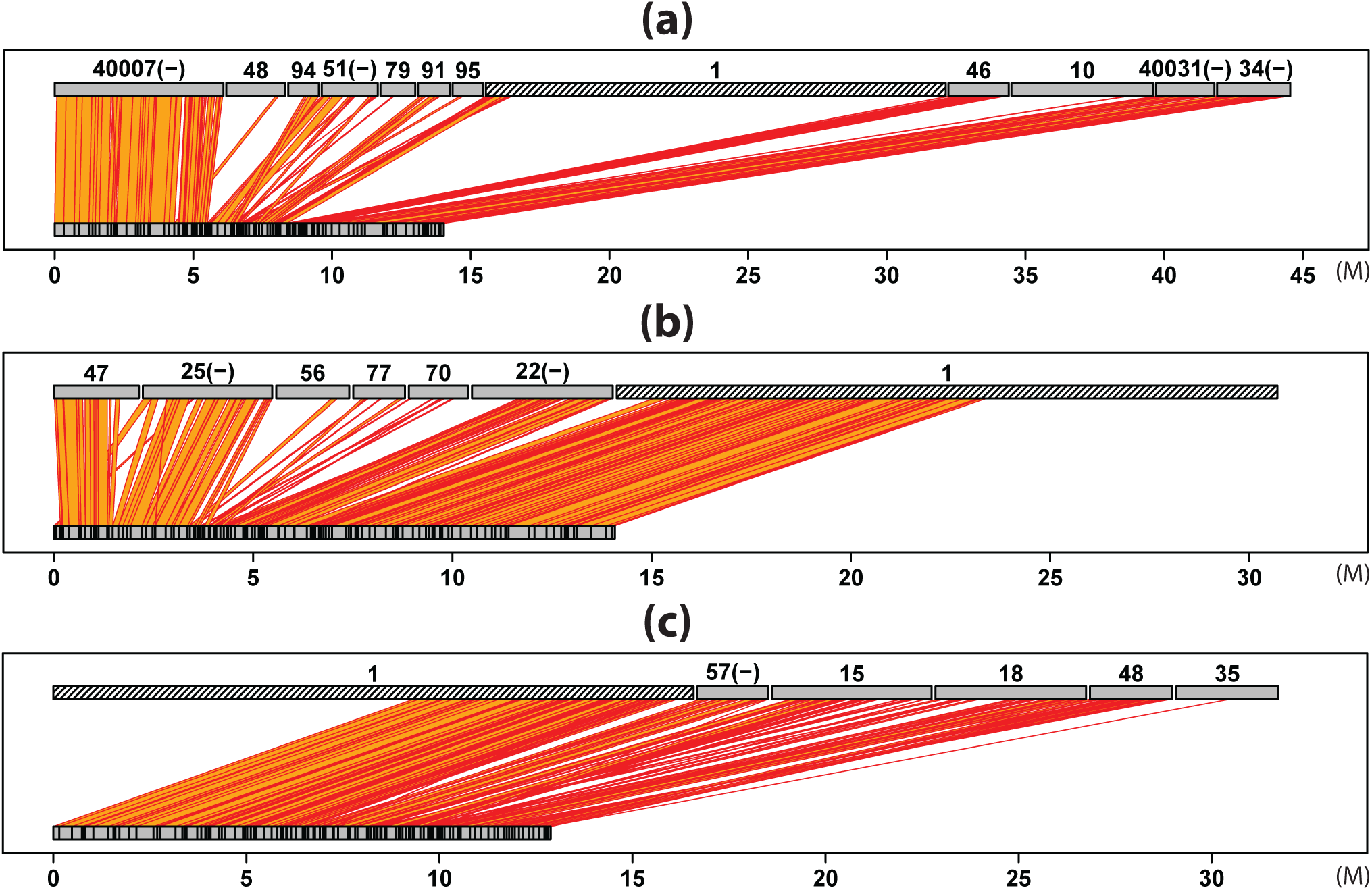
Syntenic maps of the genetic linkage map constructed from the M9xM19 GBS data and the NSP306v2 de *novo* genome assembly. (a)-(c) depict the syntenic map for three linkage groups. The scaffolds are represented by gray blocks. NSP306v2 scaffolds are placed at the top, while the ITR_r1.0 scaffolds are at the bottom. NSP306v2 scaffolds labeled with ‘(−)’ are placed in reverse direction. The apparent ‘banding’ pattern in the bottom scaffolds is just due to the fact that many of the ITR_r1.0 scaffolds are too small to distinguish by eye. A quadrilateral is used to depict the mapping information between an upper and a bottom scaffold. The blocks that represent NSP306v2 scaffold 1 are shaded with slashes.

### Application to infinium 8303 potato array data

B2721 is a tetraploid potato mapping population developed from the cross *Atlantic* × *B1829-5* consists of 156 F1 progeny. The mapping population was genotyped with Illumina 8,303 SNP array. We used the R package *fitTetra^24^* to call genotypes. The Potato Genome Sequencing Consortium (PGSC) provides an alignment of the Infinium 8303 Potato Array SNPs to the PGSC v4.03 pseudomolecules (http://solanaceae.plantbiology.msu.edu/pgsc_download.shtml). This was used to anchor the SNPs to PGSC DM v3 scaffolds^6^. The SNPs that were unaligned or aligned for multiple times were excluded from the later analysis. The scaffolds containing less than five SNPs were excluded. By using the default settings, *fitTetra* rejected 1,751 SNPs due to low *p*-value. The anchoring step further removed 1,080 SNPs. The remaining 5,472 SNPs are located on 435 scaffolds. These scaffolds summed to ~ 445Mb cover approximately 53% of the genome. We applied our method to build genetic linkage maps for these scaffolds and referred to the PGSC v4.03 pseudomolecules^6^ for validation.

The phasing algorithm failed to give any feasible haplotype calls for 126 scaffolds. This is mainly due to the low level of genetic diversity on these scaffolds. Figure 5c depicts the pairwise RF estimations against the physical distances for scaffold pairs located on the same pseudomolecules. The distances between scaffolds were estimated by the locations on the pseudomolecules. Even though there is bias compared to a conventional genetic mapping function, the estimated RFs highly correlate with the physical distances. The bias could be due to the inaccurate estimation of the physical distances between scaffolds. The gap sizes between the adjoining scaffolds on the pseudomolecules were fixed at 50Kb^6^, which is not always precise. Moreover, the pseudomolecules cover only approximately 86% of the whole genome^6^, which could also make the physical distance calculation biased. Figure 5d presents the histogram of the estimated RFs for the scaffolds from different pseudomolecules. The values are consistently large, and most of them are greater than the predefined threshold for grouping. For genetic linkage map construction, 309 scaffolds were assigned to 12 linkage groups, corresponding to the 12 PGSC v4.03 pseudomolecules, without miss assignment (Supplementary Figure S2.). The order of the scaffolds on the linkage map is consistent with that on the pseudomolecules despite a few exceptions for scaffolds that are closed to each other (Fig. 7).

**Figure 7.**
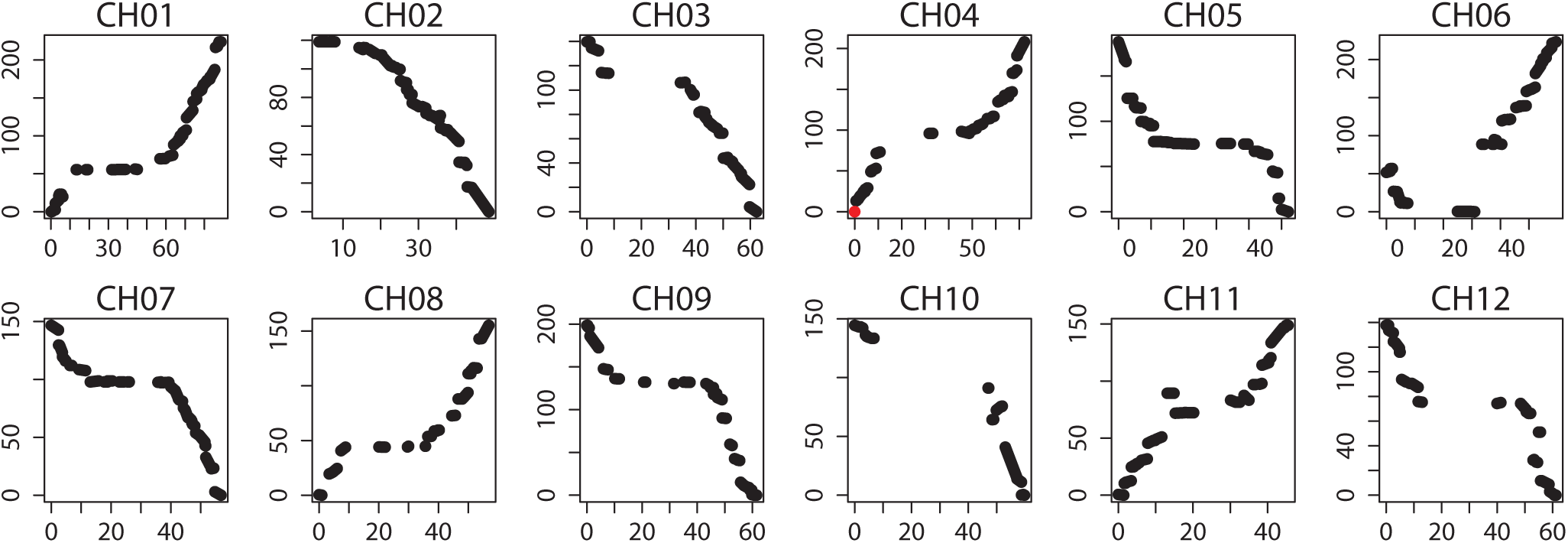
Genetic linkage map constructed from *Atlantic × B1829*-5 mapping population 8303 potato array data. *x*-axis represents the positions on the PGSC v4.03 pseudomolecules (Mb) and y-axis represents the positions on the genetic linkage map (cM).

We applied this method to another Infinium 8303 potato array data for a outcrossed family generated by *12601ab1* × *Stirling^25^*. This family consists of 192 samples. The variant calling process ended up with 3,988 SNPs from 358 scaffolds for linkage analysis. 240 scaffolds were reserved in the final genetic linkage maps. 12 linkage groups were constructed corresponding to the 12 pseudomolecules. The results are very similar to that described for the B2721 dataset (Supplementary Figure S7., Supplementary Figure S2.).

### Comparison to well-known genetic mapping tools

We compared our genetic mapping method to three existing tools designed for outcrossed population including Onemap^17^ and Lep-MAP2^19^ for diploid and TetraploidMap^20^ for tetraploid. The marker sets used for comparison consist of 1000 SNPs sampled from a simulated outcrossed family with known linkage groups and order (Online Methods). The missing data rates range from 0 to 0.9, while the genotyping error rates range from 0 to 0.5. For each configuration, we conducted 30 independent experiments for PolyGembler, Onemap, and Lep-MAP2, and two experiments for TetraploidMap as only a graphical user interface is provided by the author. Moreover, TetraploidMap requires one of the parents is nulliplex. SNPs that are not handled by TetraploidMap were removed. This process discarded ~ %50 SNPs for each dataset.

The results are reported in Table 1 with details in Supplementary Table S8. As we can see, PolyGembler is always better or at least not worse than the other methods in accuracy for grouping and consistently outperformed the other methods in accuracy

**Table 1.** Results for comparisons of *PolyGembler* to *Onemap, Lep-MAP2* and *TetraploidMap.* The statistics for *PolyGembler, Lep-MAP2* and *Onemap* are averaged on 30 independent runs (Intel^®^ Xeon^®^ Processor E5-2667 v3 CPU, 3.20GHz), while for *TetraploidMap* are averaged on 2 independent runs (Intel^®^ Core^™^ i5-2520M CPU, 2.50GHz). Column γ and η report the rates of missing data and genotyping errors for the dataset. Column *E* reports the error rates of marker order on the genetic linkage map compared to the order on the chromosomes^26^. Column *NMI* reports the normalized mutual information of the constructed linkage groups and true groups. *NMI* ranges from 0 to 1, measures the mutual dependence between two random variables^27^. Larger *NMI* values indicate that the two groupings are more similar, and specifically *NMI=1* means they are identical. Column *T* reports the CPU time in seconds. Column *N* reports the number of SNPs in the genetic linkage map. *N* is sometimes less than 1000 because SNPs are either filtered out or failed to be assigned to any linkage group. Column *N* is not reported for *PolyGembler,* as it is always 1000. *PolyGembler*(2) and *PolyGembler*(4) columns report the results for diploid and tetraploid respectively. *NA* means not applicable at corresponding settings as methods return no meaningful results. TetraploidMap constructs genetic linkage maps for two parents separately, and therefore the statistics were averaged on two parental linkage maps. for ordering. Moreover, PolyGembler is able to deal with dataset with much higher missing and error rates. Here, we call a grouping with *NMI* ≥ 0.9 acceptable. For the diploid simulation, we gain acceptable results for the dataset with 80% missing genotypes from PolyGembler, while only 50% and 40% is allowed for Onemap and Lep-MAP2, respectively. The performance on the genotyping errors shows a similar trend. The largest error rate that PolyGembler is able to deal with reaches 40%, while for Onemap and Lep-MAP2, it drops to 10% and 5%, respectively. PolyGembler also outperforms the other two methods for the data with mixtures of missing data and genotyping errors. The upper bound for PolyGembler stands (30%, 10%), while for Onemap and Lep-MAP2 is (20%, 5%). Moreover, Lep-MAP2 filtered out more than half of the SNPs. Regarding the running time, PolyGembler is the fastest, followed by Lep-MAP2, and both are much faster than OneMap. It should be noted here that sometimes Lep-MAP2 is faster than PolyGembler as it discarded many SNPs. For tetraploid dataset, TetraploidMap never gives reasonable results. In contrast, PolyGembler is able to handle data set of missing data up to 60% or genotyping errors up to 40% or the mixture of them up to (30%,10%).

| $\gamma$ | $\eta$ | PolyGembler(2) | | | OneMap | | | | Lep-MAP2 | | | | PolyGembler(4) | | | TetraploidMap | | | |
| --- | --- | --- | --- | --- | --- | --- | --- | --- | --- | --- | --- | --- | --- | --- | --- | --- | --- | --- | --- |
|  |  | *E* | *NMI* | *T(s)* | *E* | *NMI* | *T(s)* | *N* | *E* | *NMI* | *T(s)* | *N* | *E* | *NMI* | *T(s)* | *E* | *NMI* | *T(s)* | *N* |
| 0 | 0 | 0 | 1.0 | 2193 | .222 | 1.0 | 12135 | 1000 | .048 | 1.0 | 2984 | 999.9 | .001 | 1.0 | 11758 | .149 | .856 | 7564 | 504.5 |
| 0.05 | 0 | 0 | 1.0 | 2127 | .225 | 1.0 | 13407 | 1000 | .069 | 1.0 | 3424 | 902.1 | .001 | 1.0 | 11848 | .147 | .831 | 9526 | 503 |
| 0.1 | 0 | 0 | 1.0 | 2195 | .221 | 1.0 | 15974 | 1000 | .069 | .999 | 2901 | 812.1 | 0 | 1.0 | 12309 | .182 | .831 | 11499 | 503 |
| 0.2 | 0 | 0 | 1.0 | 2089 | .236 | 1.0 | 18709 | 1000 | .079 | .990 | 1966 | 641.8 | 0 | 1.0 | 13650 | .162 | .855 | 6985 | 500 |
| 0.3 | 0 | 0 | 1.0 | 2126 | .220 | 1.0 | 24835 | 1000 | .084 | .978 | 1274 | 494.4 | .001 | 1.0 | 15026 | .190 | .797 | 15848 | 500 |
| 0.4 | 0 | 0 | 1.0 | 2307 | .243 | .999 | 31397 | 1000 | .097 | .937 | 720 | 351.6 | .003 | 1.0 | 17441 | NA | NA | NA | NA |
| 0.5 | 0 | 0 | 1.0 | 2161 | .270 | .999 | 41574 | 999.3 | .127 | .876 | 274 | 239.2 | .005 | 1.0 | 20472 | NA | NA | NA | NA |
| 0.6 | 0 | 0 | 1.0 | 2319 | .263 | .826 | 20912 | 962.0 | NA | NA | NA | NA | .015 | 1.0 | 23207 | NA | NA | NA | NA |
| 0.7 | 0 | .001 | 1.0 | 2432 | .197 | .583 | 352 | 351.2 | NA | NA | NA | NA | .122 | .785 | 22184 | NA | NA | NA | NA |
| 0.8 | 0 | .017 | .996 | 2703 | NA | NA | NA | NA | NA | NA | NA | NA | .157 | .324 | 9770 | NA | NA | NA | NA |
| 0.9 | 0 | .134 | .326 | 2149 | NA | NA | NA | NA | NA | NA | NA | NA | NA | NA | NA | NA | NA | NA | NA |
| 0 | 0.01 | 0 | 1.0 | 2283 | .245 | 1.0 | 13355 | 1000 | .065 | 1.0 | 4155 | 978.2 | 0 | 1.0 | 11984 | .133 | .795 | 10452 | 515 |
| 0 | 0.02 | 0 | 1.0 | 2280 | .238 | 1.0 | 18206 | 1000 | .069 | 1.0 | 3892 | 954.5 | .001 | 1.0 | 12159 | .173 | .797 | 7241 | 519.5 |
| 0 | 0.05 | 0 | 1.0 | 2146 | .279 | 1.0 | 37039 | 1000 | .065 | .997 | 3414 | 837.3 | .001 | 1.0 | 12716 | .154 | .785 | 5983 | 525.5 |
| 0 | 0.1 | 0 | 1.0 | 2171 | .363 | 1.0 | 80396 | 1000 | .080 | .852 | 787 | 545.7 | .001 | 1.0 | 13513 | .176 | .789 | 6124 | 527.5 |
| 0 | 0.2 | .001 | 1.0 | 2243 | .333 | .682 | 11136 | 709.2 | NA | NA | NA | NA | .003 | 1.0 | 16133 | NA | NA | NA | NA |
| 0 | 0.3 | .008 | 1.0 | 2191 | NA | NA | NA | NA | NA | NA | NA | NA | .015 | 1.0 | 17452 | NA | NA | NA | NA |
| 0 | 0.4 | .063 | .908 | 2123 | NA | NA | NA | NA | NA | NA | NA | NA | .037 | .993 | 15950 | NA | NA | NA | NA |
| 0 | 0.5 | .111 | .748 | 2055 | NA | NA | NA | NA | NA | NA | NA | NA | .096 | .885 | 14601 | NA | NA | NA | NA |
| 0.05 | 0.01 | 0 | 1.0 | 2164 | .233 | 1.0 | 11963 | 1000 | .067 | 1.0 | 3488 | 882.1 | .001 | 1.0 | 12425 | .159 | .807 | 8083 | 512 |
| 0.1 | 0.02 | 0 | 1.0 | 2214 | .238 | 1.0 | 19078 | 1000 | .068 | .998 | 2705 | 770.6 | .001 | 1.0 | 13043 | .196 | .775 | 12183 | 521 |
| 0.2 | 0.05 | 0 | 1.0 | 2241 | .303 | .998 | 53778 | 1000 | .082 | .926 | 978 | 496.8 | .001 | 1.0 | 15260 | .194 | .766 | 6331 | 526.5 |
| 0.3 | 0.1 | .001 | 1.0 | 2303 | .295 | .674 | 4055 | 72.3 | NA | NA | NA | NA | .006 | 1.0 | 18849 | NA | NA | NA | NA |
| 0.4 | 0.2 | .090 | .889 | 2146 | NA | NA | NA | NA | NA | NA | NA | NA | .099 | .844 | 16874 | NA | NA | NA | NA |

## Discussion

We described a genetic mapping method that harnesses genotyping data from outcrossing mapping populations and reference genome assembly. By mapping genotyping data to assembly, we perform linkage analysis at scaffold level. Compared to conventional marker-based methods, our method is more accurate, robust and efficient. The scaffold level linkage maps can be easily converted to chromosome-scale pseudomolecules if they cover a large portion of the chromosomes. Even if a linkage map does not represent the entire chromosome due to low quality of reference assembly or genotyping data, it provides insightful information about the order of the scaffolds, which could be utilised in pseudomolecule construction or genetic analysis.

Haplotype phasing for scaffolds makes it possible to obtain accurate RF estimations even with abundant missing data and genotyping errors. Scaffold level linkage analysis effectively reduces the computational complexity. Presumably, a well-designed heuristic algorithm is required to order the markers within each linkage group as it is NP-hard. In our method, however, the size of the problem is dramatically reduced because of the scaffold level design, which enables us to use the exact TSP solver CONCORDE^28^.

The haplotype phasing algorithm is a modification of *polyHap*^29, 30^. Compared to the original model, the hidden state space is redesigned by integration of the pedigree structure. The recombinations of parental haplotypes are restricted between those haplotypes from the same parent. In contrast, recombinations are allowed between any ancestral haplotypes in *polyHap*. Recombination restriction dramatically reduces the computational complexity, and make higher ploidy haplotype phasing possible. Moreover, we extended the model to handle multipoint haplotype phasing. The multipoint analysis runs on a superscaffold built from nearest neighbor joining, which is critical to guarantee accurate haplotype phasing for short scaffolds and high ploidy genomes.

A key feature of our method is its robustness to missing data and genotyping errors. Haplotype phasing plays an important role. Joint analysis of multiple markers on the same scaffold provides more linkage information. By reconstructing haplotypes for scaffolds, the missing genotypes are actually being imputed though we do not have an explicit imputation step. Moreover, we allow a parental haplotype to pass more than one allele at a marker locus probabilistically. This is designed to deal with genotyping errors. If a genotyping error occurs, the model will choose between introducing a recombination at that position or passing an allele with lower probability, whichever maximizes the overall likelihood. Alternatively, we could require a parental haplotype to pass exactly one allele as what happens in reality. In this case, however, as long as a genotyping error occurs, an incorrect parental haplotype recombination is introduced. It should be noted here that when the genotyping error rate is low and the model converges well, the probability of passing a certain allele by a haplotype should be close to 1.

Haplotype phasing is a critical step in our method. The accuracy of haplotype phasing is influenced by several factors. First, the size of the marker set, including the size of the mapping population and the number of the markers along a scaffold. A larger sized marker set provides more correlation information between samples as well as markers, which enhances the robustness of the model to missing data and genotyping errors. Secondly, the quality of the data, including the missing values, the genotyping errors, and the level of genetic diversity. Higher missing and error rates place a higher burden on the analysis. A low level of genetic diversity can also compromise the algorithm, especially for polyploid species. As many parental haplotypes will pass the same allele at a given position, the algorithm struggles to distinguish between them, resulting in possibly biased phasing results. This is especially true when the size of marker set is small.

The generalization of the proposed method to high ploidy genomes is straightforward. In this study, we focused on diploid and tetraploid. However, this method is capable of constructing genetic linkage maps for higher ploidy genomes. Our biggest challenge when dealing with higher ploidy species is computational, especially in the haplotype phasing step. For hexaploid, the number of hidden states increases to 14400. This is a large number of states but remains computationally tractable. However, for higher levels of ploidy, computation does become difficult.

## Methods

### Simulation of Ipomoea Trifida reference chromosomes

In order to make the simulation comparable to the real genome, we incorporated the sequencing data provided in^23^. The data consists of 709,879,448 100bp long paired-end reads with expected insert size 300bp. Firstly, we built an assembly graph from the sequencing data using SOAPdenovo2^4^. Then, we extracted all the contigs and corresponding coverage from the assembly graph. We assumed the coverage represents the copy number of a contig in the reference genome. In total, 1,669,780 contigs summed to 494,234,273bp were identified. The longest contig is 41,310bp and the N50 statistic is 1,153bp. Contigs shorter than 100bp were filtered out. After filtering, 881,037 contigs summed to 481,711,030bp were left for reference genome construction. Among which, 502,551 contigs of 365,400,227bp have exactly one copy, while the remaining 378,486 contigs of 116,310,803bp have more than one copy, represents approximately 75.85% and 24.15% of the whole genome, respectively (Supplementary Table S1.). Here, for convenience, denote **P** and **P̅** the contig pool formed by the non-repeated and repeated contigs, respectively. In order to construct longer repeats, contigs were randomly selected from **P̅** and combined. We generated five categories of long repeats distinguished by size, i.e., values fall into half-closed intervals [0,500), [500,1000), [1000,10000), [10000,20000), and [20000,40000), respectively. To generate a long repeat for a given category, we randomly sampled a number *L* from the interval as the target repeat length, and randomly sampled a number *k* from 2, 3 and 4 with probabilities 0.9, 0.09 and 0.01 respectively as the copy number of this long repeat. Everytime, we randomly selected a short repeat with at least k copies from **P̅** and added it to the end of the long repeat until the length is equal to or greater than L, or there is no available short contig any more. Once a contig was used, the copy number it presents in **P̅** was subtracted by k. Once the construction of a long repeat was completed, k copies of this contig were added to **P** (Supplementary Table S2.). It should be noted here that we started from the longest category and all the way down to the shortest one to reduce the risk of running out of repeated contigs for long repeats. We assumed each of the five categories long repeats represents similar portion of the whole genome, namely ~ 23Mb. We recorded the total length of the long repeats in the current category, and proceeded to the shorter one when it exceeds the expected value. Once the long repeat construction was completed, we generated 15 chromosomes using all the sequences in **P** by random concatenation (Supplementary Table S3.).

### Simulation of reference scaffolds

ART (Version 2.5.8)^31^ was employed to simulate Illumina HiSeq 2000 reads from the reference chromosomes. Three libraries were generated including a paired-end library with insert size 300bp (standard deviation 10bp) at 120x and two mate-pair libraries with insert size 3000bp (standard deviation 50bp) and 10000bp (standard deviation 200bp) respectively at 30x. The read length was set to 100bp and all the other parameters were set as default (Supplementary Table S4.). Jellyfish (version 2.2.6)^32^ (*k* = 17,23,31) was applied to calculate the *k*-mer frequency of the sequence data. The genome size was estimated as 490,480,568bp with *k* = 17 (Supplementary Figure S1.). SOAPdenovo2 (Version 2.04)^4^ was used to assemble the genome. The paired-end reads were used to build contigs and mate-pair reads were used for scaffolding. The resulted genome assembly consists of 402,557 scaffolds summed to 526,864,946bp. The longest scaffold is 669,577bp and the N50 statistic is 106,352bp (Supplementary Table S5.).

### Simulation of marker sets used for comparison

First, we simulated two parental genotypes using the simulated reference chromosomes (Supplementary Table S3.). We only considered SNPs. Since the SNP density varies across different regions, the distances between adjacent SNPs were sampled from a mixture of *K* Poisson distributions,

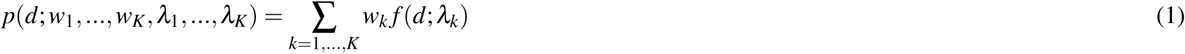

where *d* is the distance need to sample, *w =* { *w*_1_,…, *w_k_* } and *λ* = { *λ*_1_,…, *λ_K_* } are the weights and parameters of the *K* components, respectively, and *f* (⋅) is the Poisson probability mass function. We set *K* = 7, w = {0.05,0.10,0.20,0.30,0.20,0.10,0.05} and λ = {50,100,200,500,1000,2000,5000} and generated ~ 565K SNPs. Next, we simulated the meiosis process of the two parents with the software PedgreeSim V2.0^33^ to produce 190 F1 progeny genotypes. Then, to generate a marker set, we randomly selected 100 contigs of 11cM from the chromosomes, and sampled 10 SNPs from each contig. The total genetic length of the chromosomes is 1500cM (Supplementary Table S3.), and therefore the marker set covers approximately 73% (1100/1500) of the genome. Finally, we randomly introduced missing data and genotyping errors to the marker set.

### Simulation of GBS data

We mapped the genotypes generated by the software PedgreeSim V2.0^33^ back to the simulated reference chromosomes to produce genome for each sample. To generate GBS data, we simulated the GBS protocol to produce sequences with *ApeKI* as restriction enzyme^12^. Substitutions, insertions and deletions were randomly introduced as sequencing errors. The overall error rate was set to 1 × 10^−3^, from which substitutions, insertions and deletions were randomly sampled with equal probabilities. In order to make the simulation more comparable to the real data, we constructed a Markov model (*S,* π **P**_1_,…, **P**_*L*−1_) to generate quality scores. *S* represents the state space of the Markov model consists of all possible quality scores. π is a vector of size |*S*| indicates the initial probabilities of quality scores in the Markov chain. **P**_*l*_ is a |*S*| × |*S*| matrix defines the transition probabilities between quality scores at *l*th base, where 0 < *l < L* and *L* is the sequence length. All the parameters were learned from real GBS data. The sequencing depth of coverage for each chromosome follows a truncated normal distribution. The overall depth for diploid and tetraploid GBS data is 10.8× and 43.2×, respectively (Supplementary Note S1.)

### Variant detection of GBS data

Similar to the TASSEL-GBS pipeline^34^, we employ a tag-based design. The implementation consists of five major steps: (1) generate and encode tag sequences. The input for the program includes one or more FASTQ files containing GBS reads and a configuration file generated by the GBS protocol indicating the barcodes, the flowcells, the pedigree structure and so on. The program takes each sequence from the FASTQ files, removes the barcode, and replaces low quality bases with ‘N’ according to a predefined threshold. We call the processed sequence a tag sequence. The tag sequence is then encoded with a Bitset. For which, each nucleobase including ‘N’ consumes three bits. The encoded tag sequence is hashed and put into a hash table. For each tag, a mutable integer array is maintained to record the counts in each sample. During this progress, the hash table could become large and hard to maintain. Therefore, we constantly monitor the memory consumption by the program. If the usage exceeds a predefined threshold, 90% by default, the hash table is then written to the hard disk and cleared. The output is a binary file where each line records an encoded tag sequence and the counts. Tag sequences are sorted by hashcodes before written to hard disk. (2) merge tag sequences. There are more than one tag sequence binary files either because of multiple GBS FASTQ files or multiple volumes for a GBS FASTQ file due to memory limitation. The program merges them to produce a single file contains all the tag sequences. The major task is to calculate the total counts for tags displayed in multiple files. As the tag sequences are sorted, this process takes linear time. (3) decode tag sequences to generate FASTQ file. The encoded tag sequences in the binary file are decoded to generate a FASTQ file. The integer arrays describe the counts of tag sequences in each sample are written to a separate index file. (4) align to the reference assembly. The FASTQ file containing all tag sequences is mapped to the reference assembly. We use BWA-MEM^35^ (version 0.7.12-r1039) with default parameters. According to the index file, the resulted BAM file is split to produce BAM files for each sample. It should be noted here that, for tag sequences observed for multiple times, corresponding number of BAM records are written to the BAM file so as to ensure correct allele depth. (5) variant calling and filtering. We run Freebayes^36^ (version v0.9.20) with default parameters for variant calling and a customer script for filtering. We require a variant, (a) the allele number equals to two, (b) the minor allele frequency is no less than 0.1, (c) the percentage of missing data is no greater than 50%, and (d) the number of markers along a scaffold is no less than five. The program is multi-threaded for speed.

### scaffold haplotype phasing

We developed a hidden Markov model (HMM) to reconstruct the underlying inheritance pattern of the F1 mapping population, thus to infer the parental haplotypes, and which of these haplotypes have been inherited by each F1 progeny. This is a modification of the haplotype phasing algorithm *polyHap*^29, 30^. We redesigned the state space to reduce the computational complexity.

#### Notations

Assume the chromosome has Δ copies (so Δ = 2 for diploid genomes), and Δ is an even number. Assume *M* markers are observed along the scaffold we are interested in. We write a collection in the square brackets to indicate an ordered list, while in the curly brackets to indicate an unordered list. The elements in an ordered list are indexed and could be accessed by indices. We write π = [π _1_,…, π _*n*_] apermutation of the sequential number 1 to *n*. Then for any unordered list *o* = {*o*_1_,…, *o_n_*}, we write π (*o*) = [*o*π_1_,…, *o*π_*n*_] for a permutation of *o*, and Π (*o*) for the collection of all such permutations. Thus, for example, if *o* = {1,2}, there are two permutations, namely [1, 2] and [2, 1], whereas if *o* = {1,1}, there is only one permutation. We use unordered and ordered list to represent unphased and phased data, respectively.

Consider the mth marker on the scaffold, where *m* = 1,…,*M*. Write *Ĥ*_*m*_ = {*h_1_,…, h*_Δ_} for unphased paternal haplotypes, *H̆*_*m*_ = {*h*_Δ +1_,…, *h*_2Δ_} for unphased maternal haplotypes and *H̅*_*m*_ = *Ĥ* _*m*_ ∪ *H̆* _*m*_ for all the parental haplotypes. Since we do not allow haplotype transfers for parents, without loss of generality, the phased paternal and maternal haplotypes at *m*th marker can be written as 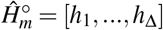 and 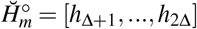, ∀*m* = 1,…, *M,* respectively. More precisely, the *δ* th copy of the paternal and maternal chromosomes at *m*th marker are denoted by 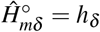 and 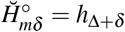 respectively, where *δ* = 1,…, Δ. The phased paternal and maternal haplotypes for the whole scaffold are then denoted by 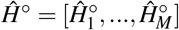 and 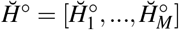, respectively. For any given F1 progeny, note here that it inherits Δ/2 haplotypes from each parent, the unphased haplotypes at mth marker can then be written as 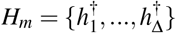, where 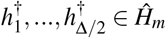 and 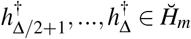. For example, in diploid species, ∪_*Hm*_ = {{*h*_1_, *h*_3_}, {*h*_1_,*h*_4_}, {*h*_2_,*h*_3_}, {*h*_2_,*h*_4_}}. Write 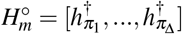 for phased haplotypes at *m*th marker, where 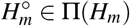, the phased haplotypes for the F1 sample is then denoted by 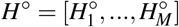.

Assume ∧_*m*_ alleles are observed at the mth marker, namely *A_m_* = {*a*_1_,…,*a*_∧*m*_}. Each allele *a*_λ_, where *λ* = 1,…, ∧_*m*_, is assumed to be descended from at least one of the parental haplotypes. Write *g_m_* = {*g*_*m*1_,…, *g*_*m*Δ_}, for the unphased genotype of a F1 sample at the *m*th marker, where g_*mδ*_ ∈*A_m_* and *δ* = 1,…, Δ. Write 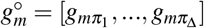 for the phased genotype, where 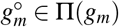, the phased genotypes for the scaffold is then written as 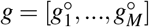.

#### Transition probability

The transitions in the HMM are designed to model the recombination events of the parental haplotypes during the meiosis process. For any F1 progeny, the probability of the δth copy of the chromosomes transfers from the parental haplotype state *h*_δ_ at (*m* − 1)th marker to *h*_τ_ at *m*th marker, where *h*_δ_, *h*_τ_ ∈ *H̅*_*m*_, is defined by,

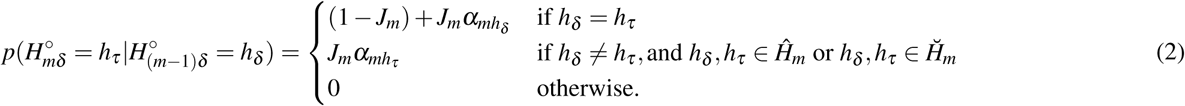

where *J_m_* is the probability of a jump occurring during the meiosis process between parental haplotypes at marker *m* − 1, and α_*mhτ*_ is the probability that this jump results in the haplotype *h_τ_* irrespective of the original haplotype. *J_m_* is positively related to the physical distance between (*m* − 1)th and *m*th marker. When the two markers are tightly linked, *J_m_* should be extremely small to prevent haplotype states jumping frequently. Based on the haplotype model, the transition probability between two phased haplotype state lists from (*m* − 1)th to *m*th marker is given by,

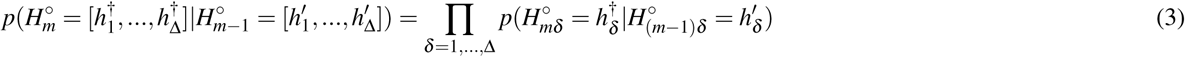

and between the unphased haplotype state lists is written as,

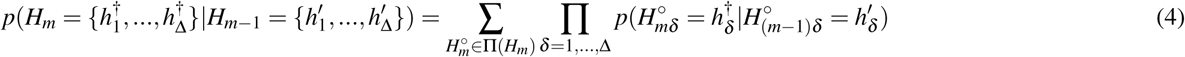

It should be noted here that, the transitions between paternal and maternal haplotypes are prohibited (see equation (2)). Equation (4) can be rewritten as,
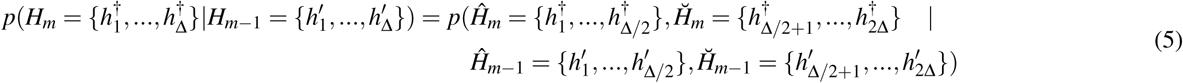

where *Ĥ_m_* and *Ĥ_m_* _− 1_ represent the paternal meiosis process, while *H̆_m_* and *H̆*_*m* − 1_ represent the maternal meiosis process. Assume the independence of the paternal and maternal meiosis,

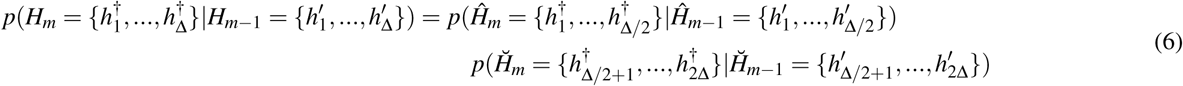

Compared to equation (4), equation (6) significantly reduces the complexity of computation.

#### Emission probability

For the *m*th marker, the unphased genotype and haplotypes of a F1 sample is written as *g_m_* = {*g_m1_*,…, *g_m_*_Δ_} and 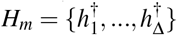, respectively. Denote *θ*_*mh*_(*a*) the emission probability of the allele *a* from the parental haplotype h at the mth marker, where *m* = 1,…, *M, a* ∈ *A_m_*, and h ∈ *H̅*. We then calculate the joint emission probability of the unphased genotype given the unphased haplotypes by,

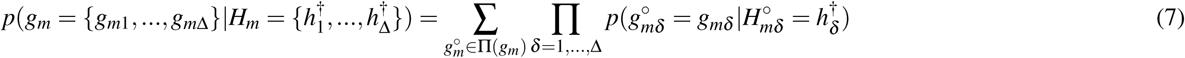

Or

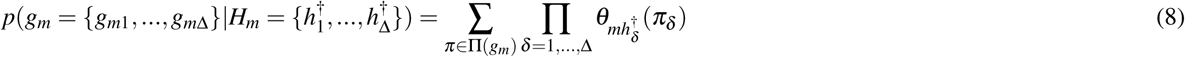

If *g_m_* is completely missing for an individual at marker m, we set *θ*_*mh*_(*a*) to be uniformly distributed over all alleles. If *g_m_* is partially missing, we set *θ*_*mh*_(*a*) to be uniform over alleles that are consistent with the observed genotype at marker *m*.

#### More computational details

We use Dirichlet priors on all parameters. Let *θ*_*mh*_ ~ *Dirichlet*(*μ_θ_m_θ_*), where *m*_θ_ is the uniform vector with each element equal to 1/Λ_*m*_, and *α*_*m*_. ~ *Dirichlet*(*μ_α_ m*_α_), where *m*_α_ is the uniform vector each element equal to 1/Δ. Also let 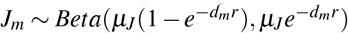, where *d_m_* is the physical distance between consecutive marker *m* − 1 and *m* along the scaffold, and *r* = 10^−8^ per base pair, reflecting the background recombination rate. We use *μ_θ_* = *μ_α_* = 1 and *μ_*J*_* = 10^5^ for initialisation of the EM algorithm and *μ_θ_* = *μ_α_* = *μ_J_* = 0.1 for the maximisation step.

The frequencies of the 2Δ parental haplotypes are expected to be equal in the mapping population. Occasionally, however, we observed huge skew in haplotype phasing results. This could be the situation when the marker set has a low level of genetic diversity. In an extreme case assume a marker set of all homozygous markers. All the parental haplotypes would then pass the same allele at a given position, thus the algorithm could not distinguish them from each other and would finally end up with biased haplotyping. Even though the homozygotes were filtered out, it still could be a problem if many of the markers lack genetic diversity especially for polyploids. Biased haplotyping were removed to avoid inaccurate RF estimations. We counted the number of each parental haplotype and require the proportion to the expected value falls into the interval 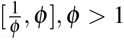. Otherwise, we discard the result for this run. The expectation of a parental haplotype in the mapping population is calculated as 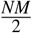 where *N* is the size of mapping population and *M* is the number of markers along the scaffold. In this study, we choose ϕ = 2.

#### RF estimation

The RF between two markers is estimated by the proportion of the number of recombinants to the total number of haplotypes in the F1 progeny. Assume a genome of ploidy Δ and a mapping population of *N* F1 progeny, and *n* recombinations are observed between the two markers in the F1 haplotypes. The RF is then calculated as,

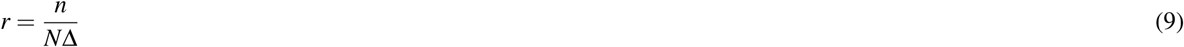

Two kinds of RFs are estimated in the proposed method, (1) in order to detect assembly errors, we calculate RFs between adjoining markers along the scaffold, and (2) in order to estimate genetic distance between two scaffolds, we calculate RFs between the outermost markers. The RF calculations within a scaffold are straightforward. We only need to count the number of jumps from one parental haplotype to another in F1 haplotypes. The RF calculations between two scaffolds, however, need to consider all four possible directions as the orientations of the scaffolds on the chromosome are unknown. Besides, as we run haplotype phasing algorithm for each scaffold separately, a parental haplotype in two phased scaffolds might be represented by different labels. As a result, we need to consider all possible correspondences, namely the permutations of the 2Δ parental haplotypes. Moreover, we also need to consider the order of the haplotypes within each F1 progeny. Therefore, 4(2Δ)! Δ ! values need to be calculated before we can decide the RF between two markers. This could be computationally impossible when the ploidy becomes large. Fortunately, we do not have to consider all these possibilities. In the HMM model, the paternal and maternal haplotypes are considered to be inherited by the F1 progeny independently, which means they can be calculated separately. The RF is then estimated as the sum of the RFs of two parents. In this way, only 8Δ!( Δ/2)! values need to be calculated, and from which we choose the minimum as the final RF estimation.

Figure 8 illustrates a simple example for the RF calculation. Here we assume a diploid genome and two scaffolds **B_1_** and **B_2_** consist of nine and seven markers, respectively. Five F1 progeny were produced from the parental cross. Figure 8a shows the haplotypes of these samples called from the HMM model. Between the third and fourth marker on **B_1_** (shaded by light gray box in Fig. 8a), a jump from parental haplotype ‘2’ to ‘1’ is observed in the last F1 progeny (shaded by dark gray box in Fig. 8a), thus the RF between the two markers is estimated as 1/10. Figure 8b demonstrates the process of RF estimation between scaffolds **B_1_** and **B_2_**. Four possible directions are calculated, namely **S_1_**-**S_2_**, **S_1_**-**E_2_**, **E_1_**-**S_2_** and **E_1_**-**E_2_**. Paternal and maternal recombinants are calculated separately. There are two possible permutations for each, and for each permutation, only one possible order. Consider calculation of the RF for the direction **S_1_**-**S_2_**, i.e., RF between marker ‘1’ and ‘a’. Firstly, the algorithm examines paternal haplotypes only, i.e., haplotypes ‘1’ and ‘2’. Assume the haplotypes for scaffolds **B_1_** and **B_2_** are the same, which means no haplotype switching. Then for the first, third and fourth F1 progeny, both marker ‘1’ and marker ‘a’ are descended from the haplotype ‘1’, so they are not recombinants, whereas the rest two F1 progeny are recombinants. Therefore the number of recombinants is counted as two. Similarly, the numbers of recombinants for the other permutations and for the maternal haplotypes are calculated. The minimum number of the sum of the paternal and maternal recombinants is two, thus the RF is estimated as 2/10 = 0.2. It is worth noting here that as long as the RFs for all four possible directions are calculated, we can easily tell the relative orientation of the two scaffolds along the chromosome. The **S_1_** end of **B_1_** and the **E_2_** end of **B_2_** is closest in this example.

**Figure 8.**
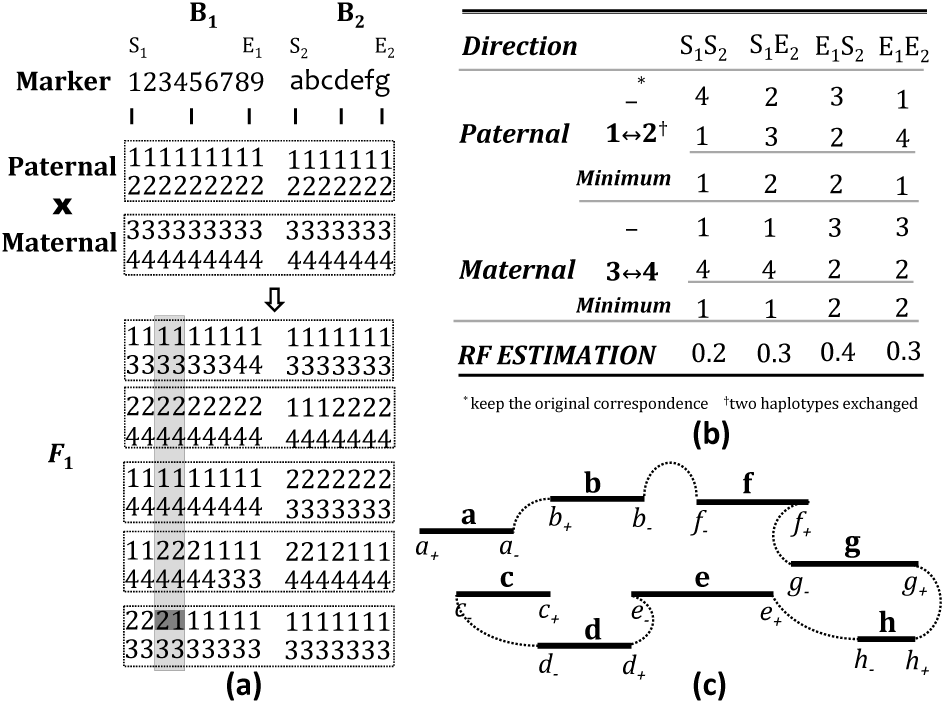
A simple example for genetic mapping. (a) the haplotype phasing results of two parents and five F1 progeny for the two scaffolds **B_1_** and **B_2_**, (b) table for RF estimation of **B_1_** and **B_2_** from the haplotype phasing results, and (c) solving TSP for a linkage group consists of scaffolds **a**-**h**.

As the Baum-Welch algorithm might be stuck in local optima, we run the haplotype phasing model multiple times with different initial parameters in order to obtain better estimations. The *K* runs with the highest likelihoods are used for RF estimation. For a pair of scaffolds, *K*^2^ RFs are calculated, and the minimum one is selected as the final estimation.

### Assembly error detection

In order to avoid the bias on the genetic linkage maps caused by the misassembly, the pipeline allows breakages of the potential incorrect scaffolds. In order to detect the misassembly positions, RFs between the contiguous markers along a scaffold are calculated. RFs are averaged on multiple runs to improve the confidence. If the RF between two adjoining markers is larger than a predefined threshold, we consider it a misassembly at that position. In this study, we set the threshold as 0.05. Apparently, the physical distance between two markers should be taken into consideration. However, it is difficult to derive a universal relationship between the physical distances and RFs, especially when the physical distance becomes large. Therefore, we do not consider positions where the distance between the adjacent markers is larger than 1Mb. The incorrect scaffolds detected by the algorithm are split at the misassembly positions to generate new scaffolds. After breakage, we run haplotpye phasing analysis on the new scaffolds and recalculate the pairwise RFs.

### Superscaffold construction and multipoint analysis

We introduce multipoint haplotype phasing analysis to improve the accuracy of RF estimations. The multipoint analysis runs the haplotype phasing algorithm on a superscaffold, which is built by joining multiple scaffolds that are close to each other according to the RFs estimated from the haplotypes. In the current study we use nearest neighbor joining to build superscaffolds. More precisely, for each scaffold, we choose the scaffold that represents the smallest RF with it, and combine them to form a scaffold pair. For each joined scaffold pair, we solve a modified TSP to calculate the order of the scaffolds thus to build a superscaffold. The gap between the two scaffolds is measured by the estimated RF. Multipoint analysis is crucial for short scaffolds as the number of markers that mapped to them could be small, which makes the haplotype phasing solely based on the marker set difficult.

### Genetic Mapping

We follow the standard framework for genetic linkage map construction. The estimated RFs are used to measure the genetic distances between scaffolds. A graph partitioning algorithm is employed to cluster the scaffolds to form linkage groups. The optimal order within each linkage group is calculated by solving a modified TSP.

#### Grouping using a graph partitioning algorithm

In order to construct linkage groups, we build a weighted graph from the pairwise RFs estimated from the haplotype phasing. The algorithm firstly constructs a symmetric *J* by *J* distance matrix denoted by *A*, where *J* is the number of scaffolds. The element at the *i*th row and *j*th column records the distance between the scaffold *i* and *j*. As we need to put more weights on the edges that connect closer scaffolds, we use *A*̅ *=* 1 − *A* as the adjacency matrix. Furthermore, in order to simplify the graph, we delete all the edges with weights less than 1 − *θ*, where *θ* is the upper bound of RF that is considered evidence for linkage. In this study, we choose *θ* = 0.38, representing a genetic distance of approximately 50cM with Kosambi mapping function. There are a plethora of graph partitioning algorithms that can be used here^37^ and the R package *’igraph’^38^* provides implementations for several of such algorithms. In the present study, we employ an information theoretic approach which detects modularity structure of a graph by minimizing expected description length of the trajectory of a random walk^39^.

#### Ordering by solving a modified TSP

We run a modified TSP to find the optimal order of the scaffolds within each linkage group. The TSP model aims to minimize the sum of the RFs along the Hamiltonian path. Figure 8c shows a simple example of eight scaffolds **a-h**. This is not a standard TSP, however, as a valid solution needs to reach a city at one end and leave from the other end. A straightforward transformation of this problem to a standard TSP is to treat the two ends of the scaffolds as different cities, which ends up with a double-sized TSP, namely, **a_+_**, **a_−_**, **b_+_**, **b_−_**, etc. The algorithm should then have mechanisms to guarantee that (1) the solution should be a Hamiltonian path instead of a Hamiltonian circuit as in a TSP, and (2) the two ends of a scaffold should be always adjacent in the final solution. The first problem is solved by simply introducing a dummy scaffold. The distances from the dummy scaffold to any other scaffolds are the same. Then we just need to cut the Hamiltonian circuit at the dummy scaffold to form a feasible solution. To solve the second problem, the distance matrix for the modified TSP is carefully designed. The distance matrix is initialised as (2*J* + 1) by (2*J* + 1) zero matrix, where *J* is the number of scaffolds. As RFs are estimated in all four directions for each pair of scaffolds, we can fill the entries represent distances between two cities from different scaffolds with the corresponding RF estimations. Now we need to fill the entries that represent distances between the two ends of the same scaffold. Assume *i* and *j* are two ends of a scaffold. We proved that *i* and *j* would be adjacent in the optimal solution if the following condition holds,

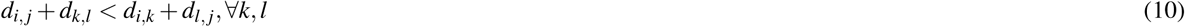

where *k* and *l* are different cities from other scaffolds, and *d_x_,_y_,* ∀ *x, y* represents the distance between city *x* and *y* (Supplementary Note S2.). As (*i*, *k*) and (*l*, *j*) are city pairs from different scaffolds, we have *d_i,k_, d_l_,_j_* ≥ 0. Therefore, equation 10 holds if,

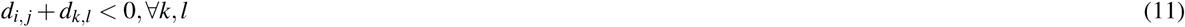

In order to guarantee equation 11, we fill all entries that represent distances between the two ends of the same scaffold with −(*φ* + ε), where *φ* represents the maximum value in the distance matrix, and ε is a small positive number. As TSP does not allow negative distances, we add *φ* + ε to every element in the distance matrix, which remains an equivalent transformation for the TSP.

Optimal solutions of the TSP are required to guarantee a feasible ordering. The TSP solver COCORDE^28^ is employed in this research. CONCORDE is an exact TSP solver which was used to obtain the optimal solution for a problem up to 85,900 cities. In our experience, it is reliable in solving the problem with the model size we usually have in the genetic mapping.

### Pseudomolecules construction

In order to improve the accuracy of the pseudomolecules, we introduce an extra step to refine the genetic linkage map. We run multipoint analysis along the entire genetic map for each linkage group and recalculate the RFs between each pair of scaffolds within this linkage group. The linkage group is then reordered by solving the modified TSP. This process is repeated for several times. The genetic map with the minimum size among the multiple runs is selected for this linkage group. This process is necessary because ordering by solving the modified TSP is very greedy, which might result in incorrect order due to just a few imprecise RF estimations. We found that this process is able to solve several incorrect ordering for the simulated data (Supplementary Figure S8.). In this study, we run this refinement process for 10 rounds.

We construct pseudomolecules according to the genetic linkage map of the scaffolds. The order of the scaffolds has been decided, thus we only need to estimate the gap size between the adjoining scaffolds. As there is a near-linear relationship between the physical distance and RF when two scaffolds are close, we calculate the average physical distance per cM represents as,

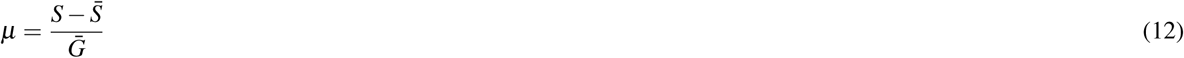

where *S* represents the genome size, and *S̅* represents total size of scaffolds been anchored in the genetic linkage maps, and *G̅* represents the sum of genetic distances between all the neighboring scaffolds. The gap between two scaffolds with genetic distance *g* is then filled with || μ*g* || ‘N’s, where |||| represents the operation to calculate the nearest integer.

### Reference scaffold true linkage groups, orders and distances

We mapped scaffolds to the reference chromosomes using the software *last* (version 73 5)^40^. A scaffold is considered being mapped to a reference chromosome if at least 50% base pairs are collinear with that and only that chromosome. The mapped scaffolds were grouped by different reference chromosomes they mapped to. The order of the scaffolds within a linkage group as well as the pairwise distances were determined by the positions on the chromosome.

There is no reference chromosome available for *Ipomoea trifida.* Instead, we used scaffolds from NSP306v2 genome assembly as references. NSP306v2 contains scaffolds longer than ITR_r1.0 scaffolds. Therefore, when we mapped ITR_r1.0 scaffolds to NSP306v2 scaffolds, multiple ITR_r1.0 scaffolds were found mapped to the same NSP306v2 scaffolds, and for those ITR_r1.0 scaffolds, according to the mapping positions, we were able to decide the order as well as the pairwise distances between them. The longest 98 NSP306v2 scaffolds (≥ 1M) were used as references. They cover approximately 58% of the whole genome assembly.

### Code availability

The software PolyGembler presented in this article and its documentation is publicly available at GitHub https://github.com/c-zhou/polyGembler.

### >Data availability

Data related to the simulation studies including the comparisons to other methods are available at the website http://data.genomicsresearch.org/Projects/polyGembler/. Data for the M9×M19 *Ipomoea trifida* and B2721 potato datasets are available upon reasonable request from Dr. Craig Yencho at. Dr. Yencho is the lead PI of the Genomic Tools for Sweetpotato (GT4SP) Improvement Project that developed the sweetpotato datasets, and he also led the development, genotyping, and phenotyping of the B2721 mapping population. Data for the *12601ab1* × *Stirling* potato dataset was provided by Dr. Christine Hackett and see^25^ for detailed data availability policy. Data related to the PGSC v4.03 pseudomolecules^6^ are available at the website http://solanaceae.plantbiology.msu.edu/pgsc_download.shtml. *Ipomoea trifida de novo* genome assembly ITR_r1.0^23^ is available at the website http://sweetpotato-garden.kazusa.or.jp. *Ipomoea trifida de novo* genome assembly NSP306v2 and the Data Release Policy is available at the website http://sweetpotato.plantbiology.msu.edu/gt4sp_download.shtml.

## Acknowledgements

We would like to thank Federico Diaz for developing M9x M19 *Ipomoea trifida* mapping population and Maria David for extracting and quantifying DNA from the M9×M19 cross. The *12601ab1*× *Stirling* infinium 8303 potato array data was provided by C.A. Hackett.

## Authors’ contributions

C.Z. and L.C. designed the study and wrote the software. D.C.G. and W.G. developed and provided the *Ipomoea trifida* mapping population materials. B.O. conducted the GBS experiment. S.W. assembled the *Ipomoea trififa* genome. C.Z. performed the analysis. C.Z. and L.C. wrote the manuscript. L.C., C.Y., A.K., M.D.C, A.G., Z.B.Z, and Z.F. supervised the project. All authors contributed to editing the final manuscript.

## Supplementary notes

**Supplementary Note S1.** More details about the simulated GBS data

**Supplementary Note S2.** Modified TSP guarantees the connection of two ends of a scaffold

## Supplementary figures

**Supplementary Figure S1.** K-mer frequency of the simulated reference genome.

**Supplementary Figure S2.** LOD score and recombination frequency heat map.

**Supplementary Figure S3.** Pseudomolecule construction for 20x tetraploid simulated GBS data.

**Supplementary Figure S4.** Potential misassembly in ITR_r1.0 scaffolds.

**Supplementary Figure S5.** Genetic linkage map constructed from Ipomoea trifida M9×M19 GBS data and ITR_r1.0 *de novo* genome assembly.

**Supplementary Figure S6.** Syntenic maps of the genetic linkage map constructed from the Ipomoea trifida M9 × M19 GBS data and the NSP306v2 *de novo* genome assembly.

**Supplementary Figure S7.** Genetic linkage map construction from Stirling × 12601ab1 mapping population SolCAP Infinium 8303 potato array data.

**Supplementary Figure S8.** Incorrect ordering of pseudomolecules solved by genetic linkage map refinement.

## Supplementary tables

**Supplementary Table S1.** Statistics of the contigs used for constructing the simulated reference genome.

**Supplementary Table S2.** Sequences used to simulate reference genome.

**Supplementary Table S3.** Simulated reference chromosomes.

**Supplementary Table S4.** Simulated HiSeq 2000 reads using ART (version 2.5.8) from the simulated reference genome.

**Supplementary Table S5.** Statistics for the simulated *de novo* genome assemly.

**Supplementary Table S6.** ITR_r1.0 potential misassembled scaffolds break points.

**Supplementary Table S7.** Misassembly identified by BioNano maps in NSP306v2 *de novo* genome assembly.

**Supplementary Table S8.** Detailed results for comparisons of *PolyGembler* to *Onemap* and *Lep-MAP2*.

## References

1. Nagarajan, N. & Pop, M. Sequence assembly demystified. Nature Reviews Genetics 14, 157–167 (2013).

2. Boetzer, M. & Pirovano, W. Sspace-longread: scaffolding bacterial draft genomes using long read sequence information. BMC bioinformatics 15, 1 (2014).

3. Burton, J. N. et al. Chromosome-scale scaffolding of de novo genome assemblies based on chromatin interactions. Nature biotechnology 31, 1119–1125 (2013).

4. Luo, R. et al. Soapdenovo2: an empirically improved memory-efficient short-read de novo assembler. GigaScience 1, 1 (2012).

5. Simpson, J. T. et al. Abyss: a parallel assembler for short read sequence data. Genome research 19, 1117–1123 (2009).

6. Consortium, P. G. S. et al. Genome sequence and analysis of the tuber crop potato. Nature 475, 189–195 (2011).

7. Ren, Y. et al. A high resolution genetic map anchoring scaffolds of the sequenced watermelon genome. PLoS One 7, e29453 (2012).

8. Howe, K. & Wood, J. M. Using optical mapping data for the improvement of vertebrate genome assemblies. GigaScience 4, 1 (2015).

9. Sequencing, T. C., Consortium, A. et al. Initial sequence of the chimpanzee genome and comparison with the human genome. Nature 437, 69–87 (2005).

10. Stone, N. E. et al. Construction of a 750-kb bacterial clone contig and restriction map in the region of human chromosome 21 containing the progressive myoclonus epilepsy gene. Genome research 6, 218–225 (1996).

11. Consortium, I. H. G. S. et al. Finishing the euchromatic sequence of the human genome. Nature 431, 931–945 (2004).

12. Elshire, R. J. et al. A robust, simple genotyping-by-sequencing (gbs) approach for high diversity species. PloS one 6, e19379 (2011).

13. Lander, E. S. & Green, P. Construction of multilocus genetic linkage maps in humans. Proceedings of the National Academy of Sciences 84, 2363–2367 (1987).

14. Manly, K. F., Cudmore Jr, R. H. & Meer, J. M. Map manager qtx, cross-platform software for genetic mapping. Mammalian Genome 12, 930–932 (2001).

15. Broman, K. W., Wu, H., Sen, S. & Churchill, G. A. R/qtl: Qtl mapping in experimental crosses. Bioinformatics 19, 889–890 (2003).

16. Iwata, H. & Ninomiya, S. Antmap: constructing genetic linkage maps using an ant colony optimization algorithm. Breeding Science 56, 371–377 (2006).

17. Margarido, G., Souza, A. & Garcia, A. Onemap: software for genetic mapping in outcrossing species. Hereditas 144, 78–79 (2007).

18. Van Ooijen, J. Multipoint maximum likelihood mapping in a full-sib family of an outbreeding species. Genetics research 93, 343–349 (2011).

19. Rastas, P., Calboli, F. C., Guo, B., Shikano, T. & Merilä, J. Construction of ultradense linkage maps with lep-map2: Stickleback f2 recombinant crosses as an example. Genome biology and evolution 8, 78–93 (2016).

20. Hackett, C. A., Milne, I., Bradshaw, J. E. & Luo, Z. Tetraploidmap for windows: Linkage map construction and qtl mapping in autotetraploid species. Journal of heredity 98, 727–729 (2007).

21. Kriegner, A., Cervantes, J. C., Burg, K., Mwanga, R. O. & Zhang, D. A genetic linkage map of sweetpotato [ipomoea batatas (l.) lam.] based on aflp markers. Molecular Breeding 11, 169–185 (2003).

22. Garcia, A. et al. Development of an integrated genetic map of a sugarcane (saccharum spp.) commercial cross, based on a maximum-likelihood approach for estimation of linkage and linkage phases. Theoretical and Applied Genetics 112, 298–314 (2006).

23. Hirakawa, H. et al. Survey of genome sequences in a wild sweet potato, ipomoea trifida (hbk) g. don. DNA Research dsv002 (2015).

24. Voorrips, R. & Gort, G. fittetra: fittetra is an r package for assigning tetraploid genotype scores. R package version 1 (2011).

25. Hackett, C. A., Bradshaw, J. E. & Bryan, G. J. Qtl mapping in autotetraploids using snp dosage information. Theoretical and Applied Genetics 127, 1885–1904 (2014).

26. Wu, Y., Bhat, P. R., Close, T. J. & Lonardi, S. Efficient and accurate construction of genetic linkage maps from the minimum spanning tree of a graph. PLoS Genet 4, e1000212 (2008).

27. Strehl, A. & Ghosh, J. Cluster ensembles - a knowledge reuse framework for combining multiple partitions. The Journal of Machine Learning Research 3, 583–617 (2003).

28. Applegate, D., Bixby, R., Chvatal, V. & Cook, W. Concorde tsp solver (2006).

29. Su, S.-Y., White, J., Balding, D. J. & Coin, L. J. Inference of haplotypic phase and missing genotypes in polyploid organisms and variable copy number genomic regions. BMC bioinformatics 9, 1 (2008).

30. Su, S.-Y. et al. Inferring combined cnv/snp haplotypes from genotype data. Bioinformatics 26, 1437–1445 (2010).

31. Huang, W., Li, L., Myers, J. R. & Marth, G. T. Art: a next-generation sequencing read simulator. Bioinformatics 28, 593–594 (2012).

32. Marcais, G. & Kingsford, C. A fast, lock-free approach for efficient parallel counting of occurrences of k-mers. Bioinformatics 27, 764–770 (2011).

33. Voorrips, R. E. & Maliepaard, C. A. The simulation of meiosis in diploid and tetraploid organisms using various genetic models. BMC bioinformatics 13, 1 (2012).

34. Glaubitz, J. C. et al. Tassel-gbs: a high capacity genotyping by sequencing analysis pipeline. PLoS One 9, e90346 (2014).

35. Li, H. & Durbin, R. Fast and accurate short read alignment with burrows-wheeler transform. Bioinformatics 25, 1754–1760 (2009).

36. Garrison, E. & Marth, G. Haplotype-based variant detection from short-read sequencing. arXiv preprint arXiv:1207.3907 (2012).

37. Buluç, A., Meyerhenke, H., Safro, I., Sanders, P. & Schulz, C. Recent advances in graph partitioning. Preprint (2013).

38. Csardi, G. & Nepusz, T. The igraph software package for complex network research. InterJournal, Complex Systems 1695, 1–9 (2006).

39. Rosvall, M. & Bergstrom, C. Maps of information flow reveal community structure in complex networks. Tech. Rep., Citeseer (2007).

40. Kielbasa, S. M., Wan, R., Sato, K., Horton, P. & Frith, M. C. Adaptive seeds tame genomic sequence comparison. Genome research 21, 487–493 (2011).

